# Harnessing Generative AI to Decode Enzyme Catalysis and Evolution for Enhanced Engineering

**DOI:** 10.1101/2023.10.10.561808

**Authors:** Wen Jun Xie, Arieh Warshel

## Abstract

Enzymes, as paramount protein catalysts, occupy a central role in fostering remarkable progress across numerous fields. However, the intricacy of sequence-function relationships continues to obscure our grasp of enzyme behaviors and curtails our capabilities in rational enzyme engineering. Generative artificial intelligence (AI), known for its proficiency in handling intricate data distributions, holds the potential to offer novel perspectives in enzyme research. By applying generative models, we could discern elusive patterns within the vast sequence space and uncover new functional enzyme sequences. This review highlights the recent advancements in employing generative AI for enzyme sequence analysis. We delve into the impact of generative AI in predicting mutation effects on enzyme fitness, activity, and stability, rationalizing the laboratory evolution of *de novo* enzymes, decoding protein sequence semantics, and its applications in enzyme engineering. Notably, the prediction of enzyme activity and stability using natural enzyme sequences serves as a vital link, indicating how enzyme catalysis shapes enzyme evolution. Overall, we foresee that the integration of generative AI into enzyme studies will remarkably enhance our knowledge of enzymes and expedite the creation of superior biocatalysts.

## Introduction

Enzymes serve as biological catalysts that expedite cellular chemical reactions without being depleted (1). They are essential in various processes, including metabolism, digestion, gene regulation, and cellular signaling. The significance of enzymes extends into biotechnology, medicine, and industry, where they find applications in the production of pharmaceuticals, biofuels, diagnostics, and targeted treatments (2, 3). Thus, improving our knowledge of enzyme and advancing in enzyme engineering is essential (4–7).

Numerous studies have focused on enzymes, particularly in exploring their structure, function, and mechanism (8–11). The conventional experimental approaches in use include enzyme kinetics analysis, X-ray crystallography, nuclear magnetic resonance spectroscopy, and cryo-electron microscopy. A bottleneck in these experimental methods, however, lies in their low-throughput, which may be augmented by computational solutions. Computational methods like molecular dynamics simulations and quantum mechanics/molecular mechanics for free energy calculations offer insights that might be otherwise challenging or highly resource-intensive with experimental approaches. However, a comprehensive and predictive understanding of enzyme behavior still presents a significant challenge.

Machine learning has recently demonstrated its capability to bolster both the analysis and engineering of proteins, including enzymes (as discussed in Refs (12–26)). Machine learning algorithms are broadly classified into two groups: discriminative models and generative models. Discriminative models specialize in data classification or labeling. In contrast, generative AI seeks to identify the innate distribution of data, thereby enabling the generation of new instances informed by this distribution. When implemented with enzyme sequences, generative models reveal latent patterns and tendencies in the immense sequence space, ultimately aiding in the identification of novel functional sequences. Several reviews have summarized the application of generative models to proteins from an engineering perspective (19, 26). There is a pressing need for an exploration aimed at deciphering proteins, particularly enzymes, in a way that resonates with biophysicists’ perspective, as they seek to decode the physical principles behind the structure and function of enzymes, guiding a more rational approach to enzyme engineering. This is the main objective of the present manuscript.

In this review, we examine the latest progress in applying generative AI to enzymes, focusing on the analysis of natural protein sequence. We begin with an overview of several notable generative models. Next, we discuss the progress made by generative models in deepening our insight into enzyme evolution, architecture, and function. In particular, generative models analyzing natural enzyme sequences could yield a metric that correlates with enzyme fitness, activity, and stability, thereby indicating how the physicochemical properties of enzymes shape their evolution. Even for *de novo* enzymes utilizing enzyme scaffolds from nature, natural evolutionary information proves to be insightful to rationalize their laboratory evolution. We then explore the application of generative models for enzyme engineering. Lastly, we discuss the challenges and future prospects of employing generative models in enzyme studies. We envision that generative AI has the potential to resolve persistent issues in enzyme research and sculpt intelligent strategies for enzyme engineering.

### An Overview of Generative Models

Generative models are a category of machine learning approach that uncovers the fundamental distribution of a dataset, enabling the creation of new samples that follow this distribution (27). This generative capability proves particularly advantageous for data synthesis tasks, such as generating text, images, or sounds resembling the original training data. Given the abundance of homologous sequences from various organisms, the application of generative models to enzyme design is a natural fit (14, 18, 19, 22, 25), as the sequence probability learned may also carry biological and evolutionary implications. In this setting, we provide a succinct introduction to several generative models pertinent to enzyme sequence studies, including maximum-entropy (MaxEnt) models (28), variational autoencoders (VAEs) (29), language models (30), and generative adversarial networks (GANs) (31).

While all these diverse models fall under the umbrella of generative AI, each offers a distinct suite of characteristics, including distinct formulations, interpretative capacities, and generative competencies, as summarized in **Table 1**. Frameworks such as MaxEnt models, VAEs, and language models explicitly furnish sequence probabilities; in contrast, GANs are not typically classified within the standard probabilistic model framework. MaxEnt models provide a more intuitive interpretation despite not explicitly addressing higher-order residue interactions as other models do. Setting them apart, language models eliminate the need for multiple sequence alignment (MSA) during training, a quality that allows them to conduct research across diverse protein families within a single model. These nuances suggest the potential for certain models to surpass others in specific tasks, highlighting the significance of deliberate model selection in machine learning research.

**Table 1.** Features of representative generative AI applied in protein sequence analysis.

| Model Type | Residue Coupling | Sequence Probability | MSA Independent | Interpretable Parameter | Alternative Name |
| --- | --- | --- | --- | --- | --- |
| MaxEnt | Pairwise | ✓ | ✗ | ✓ | DCA, EVcoupling, GREMLIN, etc. |
| VAE | Higher-order | ✓ | ✗ | ✗ | DeepSequence |
| Language | Higher-order | ✓ | ✓ | ✗ | ESM |
| GAN | Higher-order | ✗ | ✗ | ✗ |  |

#### Maximum Entropy Model

MaxEnt models constitute a specific class of probabilistic models that calculate a dataset’s probability distribution while maximizing information entropy given a set of constraints derived from the observed data (28). These models are rooted in information theory and statistical mechanics. In analyzing protein sequences, MaxEnt models usually take into consideration evolutionary conservation and pairwise residue correlation derived from MSA (**Fig 1A**). The pairwise residue correlation mirrors epistasis, wherein the impact of mutations on one residue is influenced by another residue, possibly due to residue interactions or functional associations. By integrating these constraints, MaxEnt models provide a framework for capturing the intricate dependencies and patterns present within protein sequences. MaxEnt models, which serve as a generic framework, have been referred to as Potts model, Boltzmann machine (32), DCA (33), EVcoupling (34), GREMLIN (35), CCMpred (36), among other names in protein studies.

**Fig 1.**
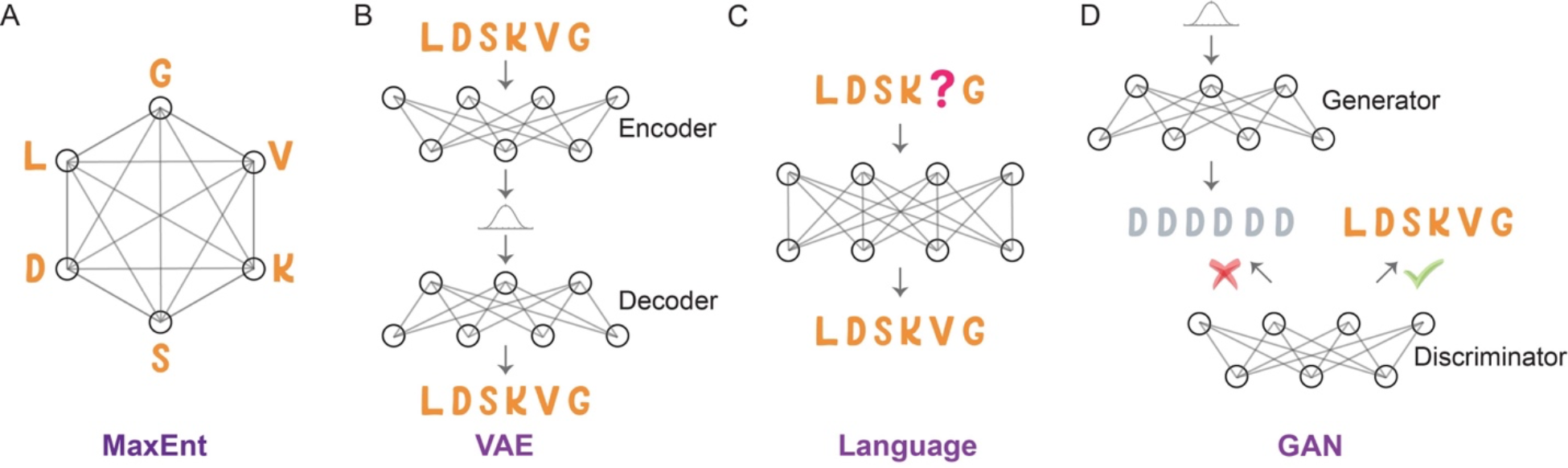
Comparative illustration of generative models utilized for protein sequence modeling. (A) MaxEnt Model: This model aims to delineate both the conservation of individual amino acids and their pairwise interactions, while concurrently making minimal assumptions by maximizing sequence information entropy. (B) VAE: A neural network that learns to encode data into a lower-dimensional latent space and then decode it back; after training, it can effectively generate new data that resembles the training set. (C) Language model: Masked language model employs a prediction-based mechanism that strives to accurately forecast the masked amino acid, thus learning the distribution of a corpus of protein sequences. (D) GAN: A framework utilizes two neural networks operating in tandem—a generator that creates new protein sequences, and a discriminator that evaluates them for authenticity.

MaxEnt models use the formula *P*(***S***) ∝ *exp*(−*E*(***S***)) to compute the probability *P*(***S***) for a sequence ***S***, where *E*(***S***) = ∑_*i*_ *h*_*i*_*S*_*i*_ + ∑_*i>j*_ *J*_*ij*_*S*_*i*_*S*_*j*_ represents the sequence statistical energy. The model parameters *h*_*i*_ and *J*_*ij*_ can be effectively trained using gradient-based optimization techniques. Depending on the conventions used, some studies might represent *E*(***S***) with a negative sign, while others might not.

#### Variational Autoencoder

VAEs are generative models that blend probabilistic modeling with deep neural network-based encoders and decoders (29). VAEs operate on the principle of mapping input data to a low-dimensional latent space. Once the encoder has successfully mapped the input data to the latent space, the decoder learns to reconstruct the original input data from this space (**Fig 1B**). The latent space provides a representation of the data’s structures. A key advantage of VAEs over traditional autoencoders is their generative ability. To achieve this, the decoder generates new data points using samples taken from the latent space. A notable instance of VAE being used on protein fitness prediction is DeepSequence (37).

The log probability of a protein sequence ***S***, log *P*(***S***), can be approximated using the evidence lower bound *E*_*q*_[log *p*(***S***|***z***)] − *D*_KL_[*q*(***z***|***S***)||*p*(***z***)] (29). In this expression, *q*(***z***|***S***) and *p*(***S***|***z***) represent neural networks responsible for encoding protein sequence data into latent variables and decoding from the latent space to reconstruct the original sequence data, respectively. The latent variables’ prior distribution, *p*(***z***), is commonly represented by a Gaussian normal distribution; the Kullback-Leibler (KL) divergence is a measure of the difference between two probability distributions.

#### Language Model

Language models used in natural language processing determine the probability distribution of a sequence of words by assigning probabilities to each potential word combination, reflecting their likelihood of occurrence within the language (30). Similarly, protein sequences can be treated as a form of language where amino acids replace words in natural languages as tokens. When tailored appropriately, protein language models can capture the distribution, patterns, and relationships observed within protein sequence data, enabling them to generate novel sequences and predict protein attributes. ESM is a widely used pre-trained language model for proteins (38).

Deep learning techniques, such as recurrent neural networks (RNNs), transformers, and autoregressive models, can be used to train protein language models. Both RNNs and transformers can be trained using masked language models, which predict a hidden amino acid based on the context provided by adjacent amino acids in the sequence (**Fig 1C**). The probability of a protein sequence ***S*** is given by *P*(***S***) = ∏_*i*_ *p*(*S*_*i*_|*S*_−*i*_). In the case of autoregressive models, they are employed to model protein sequences by learning the probability distribution of amino acids in a sequence, with each amino acid conditioned on its predecessors. The probability of a protein sequence ***S*** can be computed using the chain rule of probability *P*(***S***) = ∏_*i*_ *p*(*S*_*i*_|*S*_*1*,_ … *1 S*_*i*−1_).

#### Generative Adversarial Network

GANs comprise two distinct neural networks: the generator and the discriminator (**Fig 1D**) (31). The generator network takes random noise as input and produces new data points intended to mimic the training data. Meanwhile, the discriminator network uses both the training data and generated data as inputs to differentiate them. The training process involves refining the generator network to create data points that can deceive the discriminator network, while simultaneously improving the discriminator network’s ability to accurately discern between real and fake data points. Therefore, when used in analyzing protein sequences, GANs may distinguish real protein sequences and sequences that do not follow the rules that proteins should have.

In the realm of protein studies, GANs can differentiate authentic protein sequences from those not adhering to conventional protein structures. However, unlike other generative models, GANs do not present an explicit probability distribution for their samples. For tasks that demand an estimation of sequence probability distribution, which we will delve into further below, one might lean towards models like MaxEnt models, VAEs or language models.

### Predicting Mutation Effects on Enzyme Fitness

Generative models can assign a probability to any specific enzyme sequence based on the model’s comprehension of the inherent distribution within natural protein sequence data. It is intriguing to explore the factors that contribute to the prevalence of specific sequences or variants in nature. To this end, it is essential to bridge a connection between the probability of an enzyme sequence and its phenotype as assessed in an experiment.

Deep mutational scanning (DMS) offers a high-throughput approach for measuring the functional impact of various protein variants in a single experiment, by creating a large library of protein mutants and examining their effects on a specific phenotype (39). Given that enzymes play a pivotal role in numerous biological pathways, the growth rate of a species could serve as a phenotype to gauge the impact of mutations. Generative models offer an evolutionary landscape quantifies the sequence probability. Hence, drawing correlations between sequence probability and DMS outcomes can aid in unearthing the biological implications embedded within generative models (**Fig 2A**).

**Fig 2.**
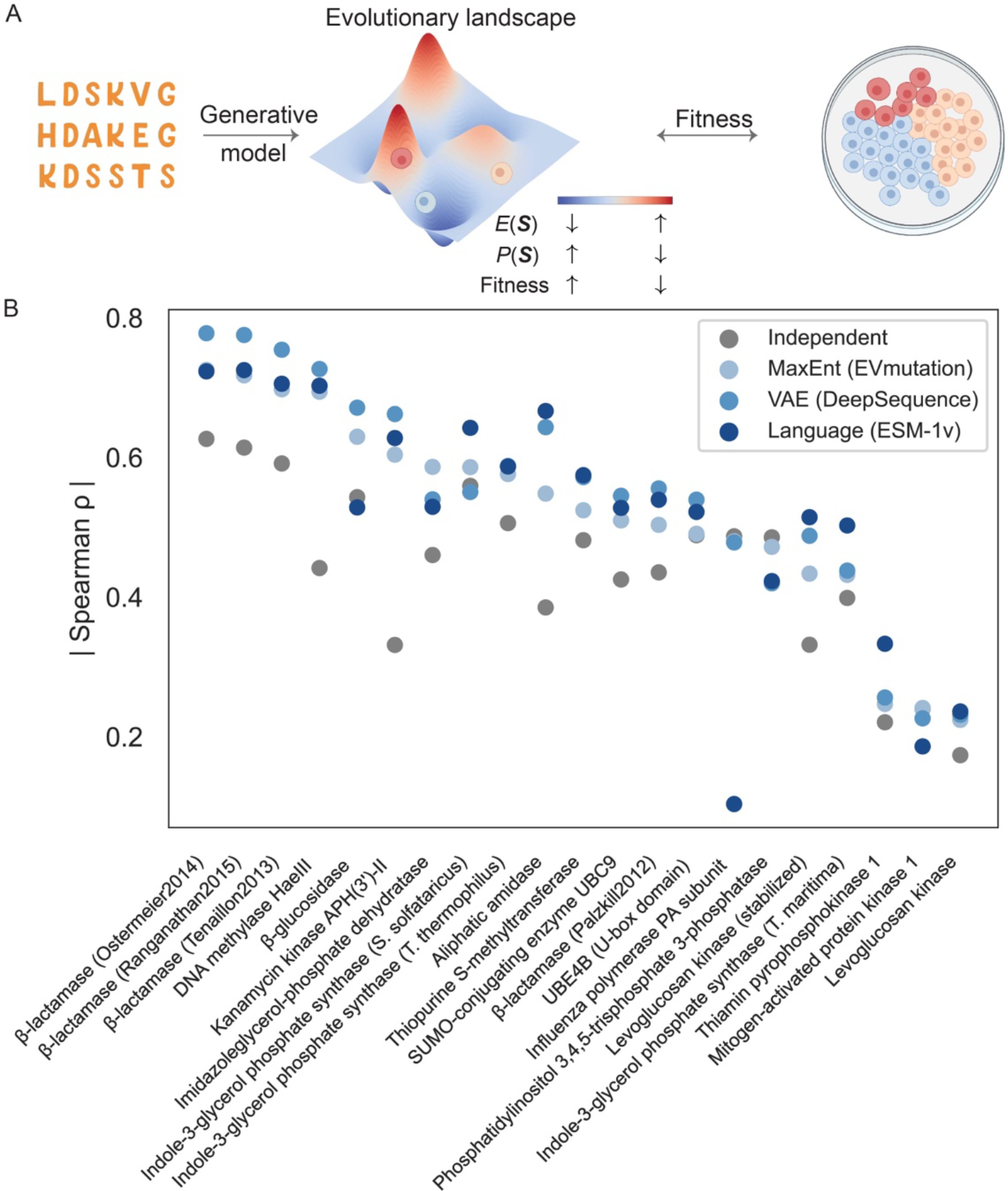
Exploration of the enzyme fitness through generative models. (A) Generative models trained on protein sequence data produce a probability distribution that correlates with protein fitness, as determined in DMS. While the evolutionary landscape is intrinsically high-dimensional, we offer a schematic illustration for explanation. Cells in the plate are illustrated using BioRender. (B) Strong correlations are evident across various generative models for a range of enzymes, as highlighted by the Spearman ρ correlation between sequence probability and fitness. The illustration incorporates several models, including the MaxEnt model, a VAE, and a language model, as represented by EVmutation (25), DeepSequence (37), and ESM-1v (38), respectively. (The data used to generate this figure were sourced from Ref (38), focusing exclusively on enzymes.)

To achieve this, several generative models have been deployed across diverse enzymes, revealing a significant correlation between sequence probability and fitness (**Fig 2B**). Overall, generative models surpass the independent model. Notably, their performance is not uniform across all enzymes. For certain enzymes, the correlations are not significant, potentially due to DMS experimental noise or the intricacies of evolution. In the following, we will showcase the differing performances of various models.

To our knowledge, the MaxEnt model was the first to associate its sequence statistical energy, or probability, with the DMS experiment. Figliuzzi et al. used DCA to investigate beta-lactamase TEM-1, an enzyme conferring resistance to beta-lactam antibiotics (40). They discovered a significant correlation between the statistical energy *E*(***S***) (i.e., log *P*(***S***)) and the minimum inhibitory concentration of the antibiotic, which assesses the enzyme’s fitness in promoting survival in antibiotic environments. In a more extensive analysis, Hopf et al. corroborated that this relationship prevails across a diverse array of enzymes and other proteins using EVcoupling (25). This correlation surpasses other metrics, such as the position specific scoring matrix (PSSM) or the conservation score derived from the MSA profile, highlighting the role of epistasis in shaping protein function.

It is still noteworthy to acknowledge that the PSSM often serves as a suitable initial approach to predict fitness in the current dataset **(Fig 2B)**. The introduction of epistasis appears to outperform the PSSM, yet in numerous instances, the enhancement brought about by the epistasis model is modest. This could be attributed to the dominance of single mutations in the DMS dataset. Although coevolutionary information stemming from the epistasis between two residues is vital for protein structure prediction (41), residue conservation seems to play an important role in predicting protein function. This aligns with the observation that, predominantly, the cumulative changes in reaction free energy resulting from single mutations closely approximate the free energy alteration measured in the multi-mutant across several proteins (42).

While the MaxEnt model generally considers second-order epistasis, explicitly incorporating higher-order interactions is difficult due to the vast number of model parameters involved. In contrast, VAEs inherently incorporate higher-order correlations. It would be intriguing to investigate the significance of higher-order epistasis in predicting fitness. Riesselman et al. employed DeepSequence, a VAE framework, to relate sequence probability (estimated as ELBO) to fitness scores, resulting in enhanced correlation values compared to the epistatic MaxEnt model for a dataset encompassing numerous enzymes and other proteins (**Fig 2B**) (37).

MaxEnt models and VAEs necessitate the creation of aligned MSA. In contrast, language models offer an added degree of flexibility, capable of employing unaligned protein sequences as input. This trait permits the inclusion of a variety of protein classes into the language model. Language models have been deployed to predict the effects of mutations in proteins, a significant proportion of which are enzymes, demonstrating comparable correlation values to other models. For instance, ESM-1v (38), trained with masked language models on UniRef90 database (98 million sequences), demonstrates an impressive capacity to predict mutation effects on enzyme fitness, marginally outperforming both the MaxEnt model and VAE (**Fig 2B**).

Despite the comprehensive dataset generated by DMS, it primarily focuses on a single functional readout, which could not capture the full complexity of a protein’s sequence-function relationship (43). This issue has been extensively discussed in the widely recognized DMS dataset of beta-lactamase TEM-1 by Firnberg and co-workers (44). Synonymous mutations may lead to discernible variations in fitness values, gauged by the minimum inhibitory concentration of the antibiotic, complicating the correlation between fitness values and changes in protein sequences. Furthermore, numerous factors influence an enzyme’s fitness, such as enzyme activity and protein abundance which could be affected by protein stability. Associating fitness with the activity and stability of enzymes remains challenging. Thus, there remains a considerable gap in using such models to further predict enzyme physicochemical properties.

In addition to the zero-shot fitness prediction illustrated above in this section, the hidden states of RNNs can be used to train supervised models for many downstream applications. For example, Alley et al. employed a RNN with the UniRef50 database (24 million sequences) as input, finding that the RNN’s hidden state could effectively predict protein stability and fitness data (45). Rives et al. also applied a transformer model to analyze the UniParc database (250 million sequences) and predict mutation effects on fitness (46). Besides, there are many noteworthy studies focused on learning protein representations that can be fine-tuned for diverse tasks (47–49).

### Predicting Mutation Effects on Enzyme Activity and Stability

Assessing enzyme activity and stability is crucial for understanding biochemical mechanisms, optimizing industrial applications, guiding therapeutic interventions, and ensuring the efficacy of enzyme-based products; however, predicting the influence of mutations on such physicochemical attributes of enzymes has proven to be particularly arduous. Compared to protein thermostability prediction (50), forecasting enzyme activity poses a more substantial challenge. It frequently necessitates multiscale simulations for modeling chemical reactions, which consume significant computational resources (4, 8). For certain mutations, the difference from the wild-type could be relatively small; for instance, a tenfold rate difference translates to a reaction barrier difference of just 1.4 kcal/mol. Such a negligible difference might exceed the precision of available physical models. Meanwhile, some mutations might provoke conformational changes (51), which add to the already convoluted task of modeling enzyme catalysis.

Aside from physics-based models, predicting the effects of mutations on enzyme activity using supervised learning is currently unfeasible, largely because of the limited data on enzyme turnover numbers (52). Although millions of enzyme mutations have been measured and cataloged in databases such as BRENDA and SABIO (53, 54), many entries either lack information on experimental conditions or were measured under varying conditions in different laboratories, thus complicating direct comparisons. It is well understood that experimental factors like pH and temperature could considerably impact enzyme catalysis. Thus, predicting the impact of mutations on enzyme activity continues to be both a formidable and vital undertaking that demands innovative strategies.

One might question if a connection exists between sequence probabilities in generative models and enzyme properties such as catalytic activity and thermostability. Identifying a reliable link could lead to predictions of these physicochemical traits in a manner analogous to fitness predictions. Yet, it is crucial to remember that factors determining organism fitness often interrelate. The complex interplay between enzyme catalytic activity (*k*_)*+_) and protein thermostability serves as a prime example (55–59). In light of these complexities, it is understandable why drawing a direct association between sequence probability and individual enzyme properties can be intricate.

Nevertheless, using the MaxEnt model, we scrutinized 12 representative enzyme-substrate pairs frequently examined in computational chemistry, and uncovered a link between the statistical energy *E*(***S***) and enzyme properties (60). The key is the separate consideration of distinct enzyme regions. An illustration of the relationship is depicted in **Fig 3** for the enzyme dihydrofolate reductase (DHFR). When mutations occur on residues proximal to the substrate or the active center, their sequence probabilities were found to align with enzyme activity (**Fig 3A**). In contrast, mutations on the enzyme’s scaffold, distant from the substrate, exhibited sequence probabilities that aligned more with enzyme thermostability than with enzyme activity (**Fig 3B-C**). Beyond DHFR, this observation appears consistent across enzymes that catalyze different types of chemical reactions (60).

**Fig 3.**
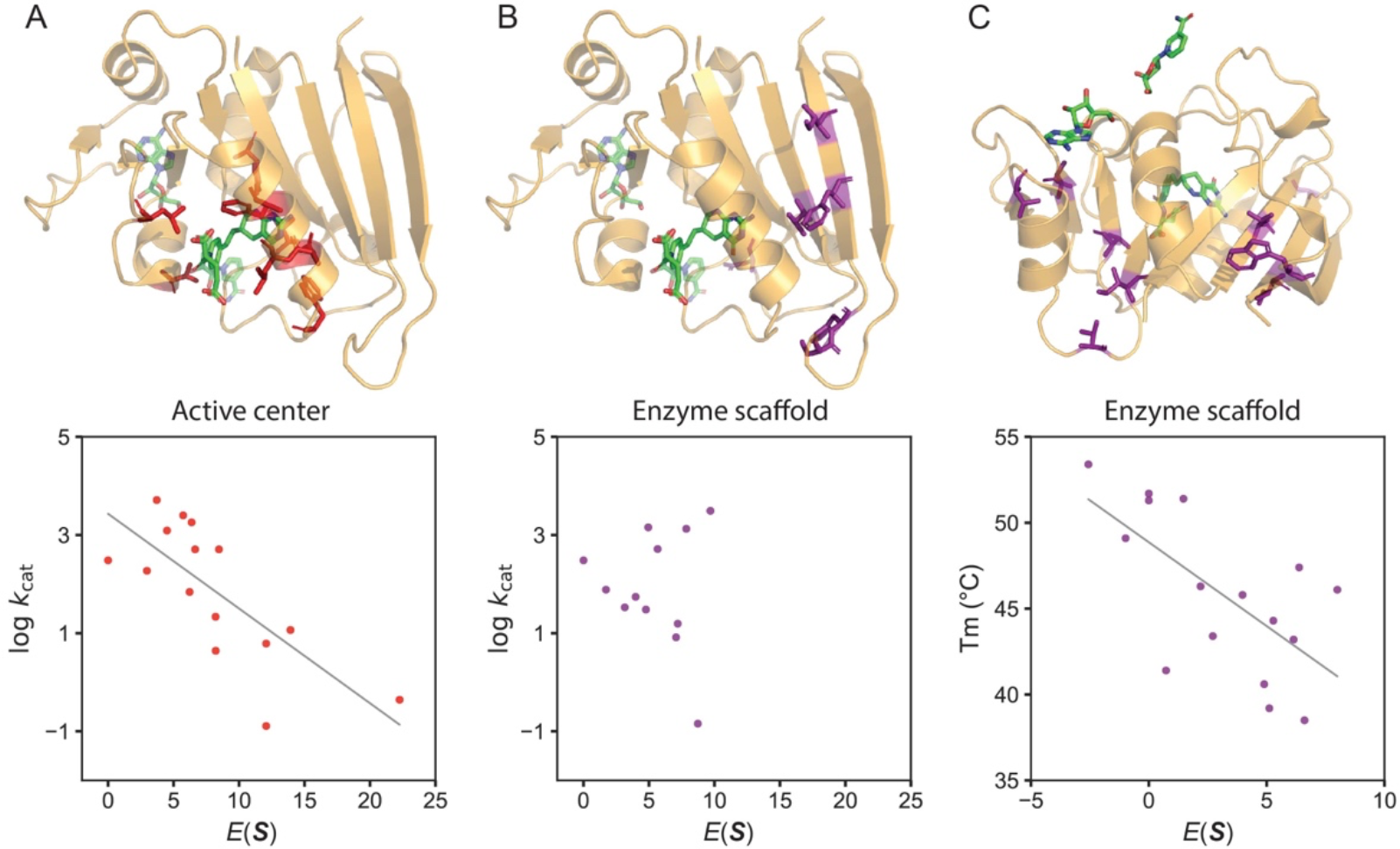
Strong correlation between enzyme physicochemical properties and evolutionary information as extracted from sequence data using generative AI, showcased using DHFR. A) The statistical energy *E*(***S***) exhibits a correlation with enzyme activity (*k*_cat_) for mutations in the vicinity of the substrate or cofactor, indicated by a Pearson’s correlation coefficient of -0.74. Mutated sites within the dataset are highlighted in red. (B) The *E*(***S***) does not show any correlation with enzyme activity for mutations located on the enzyme scaffold. Mutated sites within the dataset are denoted in purple. (C) For mutations occurring on the enzyme scaffold, the *E*(***S***) displays a correlation with thermostability (*T*_*1*_), reflected by a Pearson’s correlation value of -0.65. Mutated sites are also marked in purple. (The data used to generate this figure were sourced from Ref (60).)

It is known that structural and functional constraints influence sequence variation rates at diverse sites (61, 62), and the study of patterns within existing sequences offers valuable perspectives on protein functionality (63). Here the generative model serves as a structured framework for extracting otherwise elusive evolutionary information from the high-dimensional sequence landscape.

We term the identified correlations as the “evolution-catalysis relationship,” linking evolutionary information to factors that impact enzyme catalysis, including activity and stability (**Fig 4**). The relationship implies that nature predominantly utilizes different regions on enzymes to enhance different properties, thereby optimizing overall enzyme catalysis. Thus, the relationship provides a way to understand enzyme functional architecture. This comprehension becomes increasingly significant as it produces testable hypotheses. The most direct of these pertains to suggesting mutations to enhance specific enzyme properties, as demonstrated successfully in Ref (64) (also see discussions in “Applications in Enzyme Engineering”). However, given the vast diversity of enzymes, as demonstrated by the wide array of chemical reactions they catalyze, it is possible that further study of enzymes or specific regions may reveal additional categories (65).

**Fig 4.**
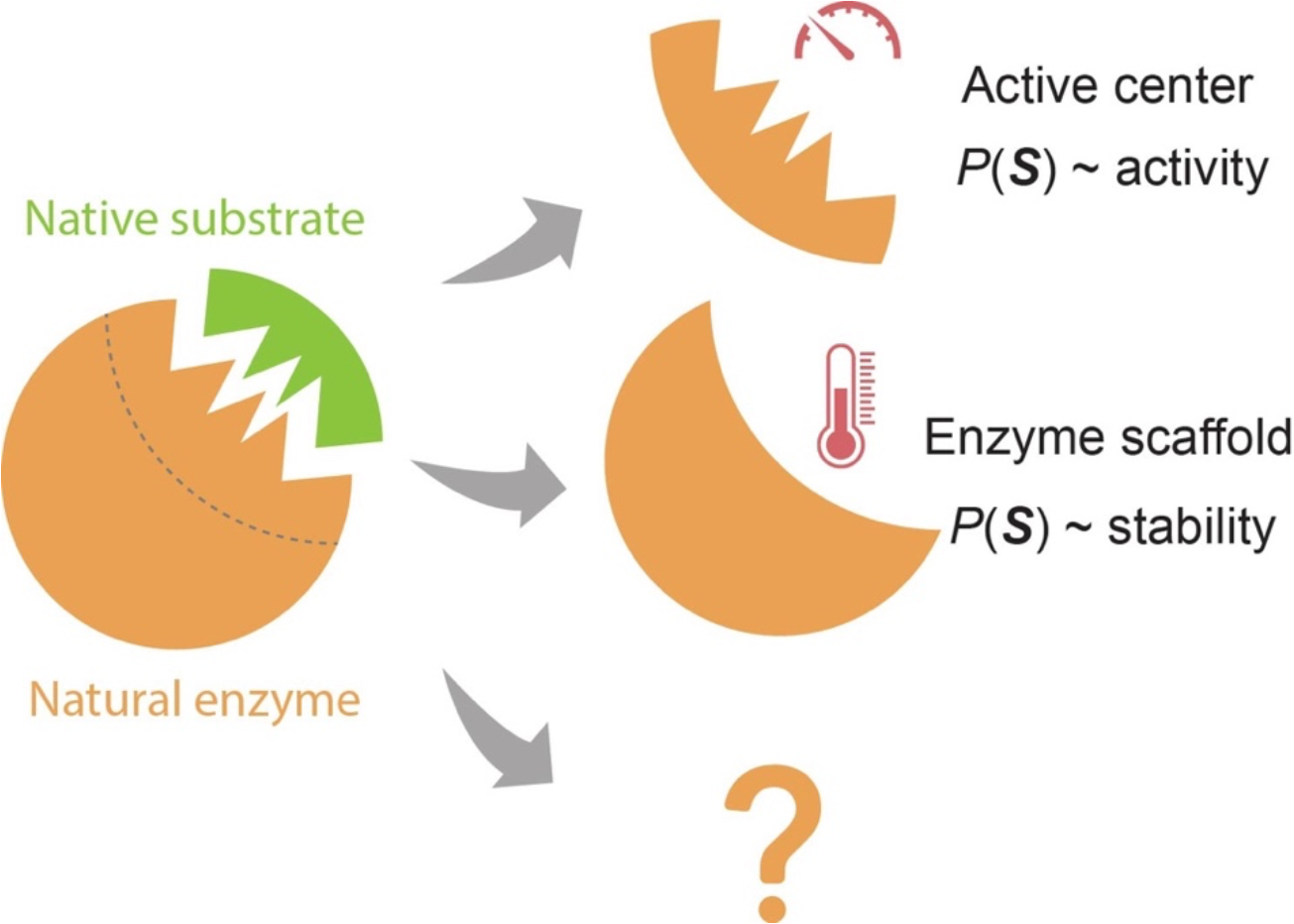
The evolution-catalysis relationship for enzyme. The sequence probability, *P*(*S*), derived from the generative model, mainly correlates with enzyme activity when mutations occur at the active center and with stability when mutations are in the scaffold region. Further examination of more evolutionary pressures and enzyme regions might reveal new classifications or fine-tune the existing ones.

It is important to clarify that not every mutation in the active center will preserve stability, and not every mutation in the enzyme scaffold will maintain enzyme activity. The evolution-catalysis relationship simply suggests that, in specific enzyme regions, some properties can be more reliably predicted using evolutionary information, while predictions for other properties might not be as certain.

A fundamental question in enzyme evolution revolves around whether natural enzymes are evolving towards higher *k*_cat_ or *k*_cat_/*K*_M_ values. While on one side, it intuitively seems that a more proficient enzyme would save an organism’s energy, it is also known that the catalytic efficiency of many enzymes is far below the diffusion limit. This paradox remains an unresolved issue, as extensively discussed by Milo and Tawfik (66, 67). However, the evident link between sequence probability and both *k*_cat_ and *k*_cat_/*K*_M_, as depicted in Fig 3A and Ref (60), implies a trend where many enzymes might indeed be on a trajectory towards enhanced catalytic efficiency during natural evolution. Without this trend, mutations that elevate enzyme activity would not be more prevalent compared to those causing reduced activity in extant enzyme sequences, and the sequence-activity correlations identified by generative AI would not be discernible. Besides, the observed correlation between evolutionary information and enzyme stability in the scaffold region aligns with studies on non-enzyme proteins (68), where stability could be the dominant evolutionary pressure.

### Rationalizing the Laboratory Evolution of *de novo* Enzymes

We expanded our research to include *de novo* enzymes, specifically focusing on Kemp eliminases, using generative models (65). In nature, there are no recognized enzymes that can catalyze Kemp elimination (69), which initially poses a challenge in leveraging natural sequence data to understand Kemp eliminases. However, we discovered that the scaffold of the Kemp eliminase, which originates from an imidazole glycerol phosphate synthase with a TIM-barrel scaffold, provides crucial insights into the function of this *de novo* enzyme (**Fig 5A**).

**Fig 5.**
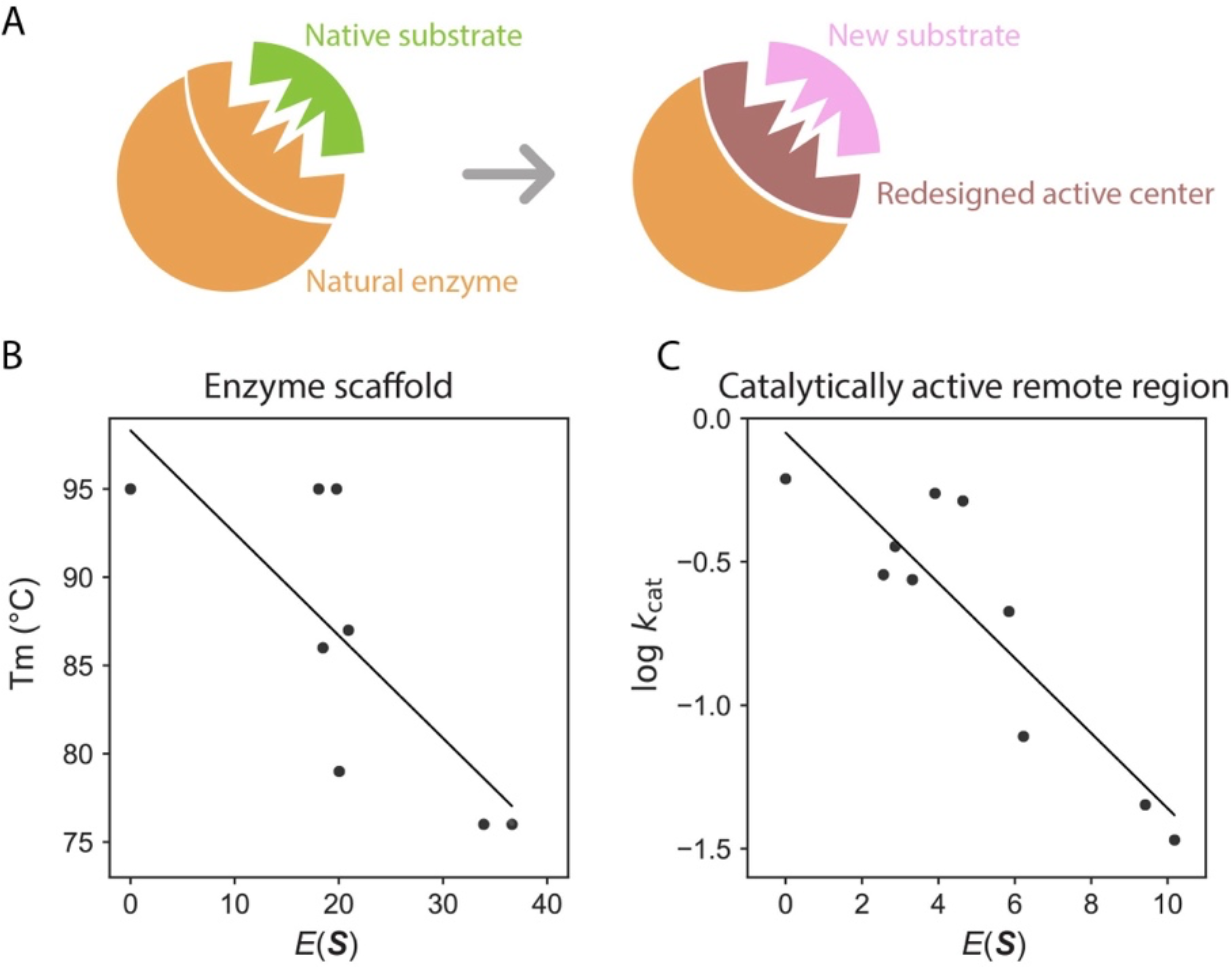
Insights from the MaxEnt model applied to *de novo* Kemp eliminase. (A) Kemp eliminase adopts the TIM-barrel scaffold of a natural synthase; its active center is modified to accommodate a new substrate. (B) Mutations on the Kemp eliminase scaffold show a correlation between *E*(***S***) and stability, evidenced by a Pearson’s correlation value of -0.77. (C) Mutations in the catalytically active remote region correlate their *E*(***S***) values with enzyme activity, indicated by a Pearson’s correlation value of -0.89. (The data used to generate this figure were sourced from Ref (65).)

We trained the MaxEnt model using homologs of the natural synthase. The mutations from directed evolution of Kemp eliminase predominantly located on the synthase scaffold; we assign each mutant a statistical energy. Notably, the statistical energy *E*(***S***) correlates with the thermostability of the Kemp eliminase variants (**Fig 5B**), although the model is trained with natural synthase homologs (65). This finding aligns with our observations regarding natural enzymes, where evolutionary information of the enzyme scaffold mainly correlates with enzyme stability (60).

There is another dataset that utilized molecular dynamics simulations and biochemistry experiments to pinpoint regions in Kemp eliminase with dynamics coupled to the active site (70). Many of these mutations were situated on the TIM-barrel loops, which we have designated as the “catalytically active remote region.” Intriguingly, mutations occurring on these regions exhibit a strong correlation between *E*(***S***) and enzyme catalytic activity, which markedly differs from other sites on the enzyme scaffold (**Fig 5C**) (65). A possible molecular mechanism could involve substrate-gated loop rearrangement determining enzyme activity, as others have posited (71). Once the substrate enters the pocket, loop rearrangement excludes water molecules from the pocket, subsequently enhancing catalytic activity. This reasoning suggests that the role of some TIM barrel loops is not overly sensitive to a specific chemical reaction, a hypothesis that could be investigated in future experiments. This might also explain the prominence of the TIM-barrel scaffold in about 10% of all enzymes.

The findings here for Kemp eliminases suggest that traits from naturally evolved enzyme scaffolds can be integrated into novel functions, illuminating the origins of new enzymatic activities.

Altogether, our studies of enzymes, encompassing both natural and *de novo* enzymes, brings fresh insights into enzyme architecture, catalysis, and evolution, giving rise to several hypotheses that warrant further investigation through biochemistry experiments and computation using physics-based models. Meanwhile, we recognize that the correlations depicted in **Fig 3** and detailed in Ref

(60) and (65) are not perfect, which is anticipated given the intertwined factors influencing enzyme evolution. Delving deeper into various evolutionary pressures and examining different enzyme regions will be crucial to further elucidate and understand the evolution-catalysis relationship.

### Decoding Protein Sequence Semantics

Generative models excel not only in predicting the consequences of mutations but also in discerning the inherent semantics and biological structures ingrained in protein sequences. The term “semantics” refers to the meaningful patterns or relationships that exist within the data - in this case, the functional and structural significance encoded in protein sequences. Here, we present several insightful studies that harness generative models to enhance our comprehension of proteins. While these investigations do not explicitly focus on enzymes, nor have they been applied to enzyme studies yet, they furnish adaptable strategies that could be customized for enzyme research in forthcoming pursuits.

Language models can learn semantics by analyzing the co-occurrence patterns of amino acids in large volumes of sequence data, or by implicitly modeling the syntax and structure of protein language. The RNN model’s hidden state was found to encode structural, evolutionary, and functional information (45). Remarkably, when projected into low-dimensional space, the embedding vectors for distinct amino acids arrange according to their physicochemical properties (**Fig 6A**). The representation of protein sequences organizes based on the organism origin and protein secondary structure (45). By carefully analyzing the correlation between neurons in the network and various properties, it is possible to comprehend the significance of different neurons. These observations extend to the transformer model as well (46).

**Fig 6.**
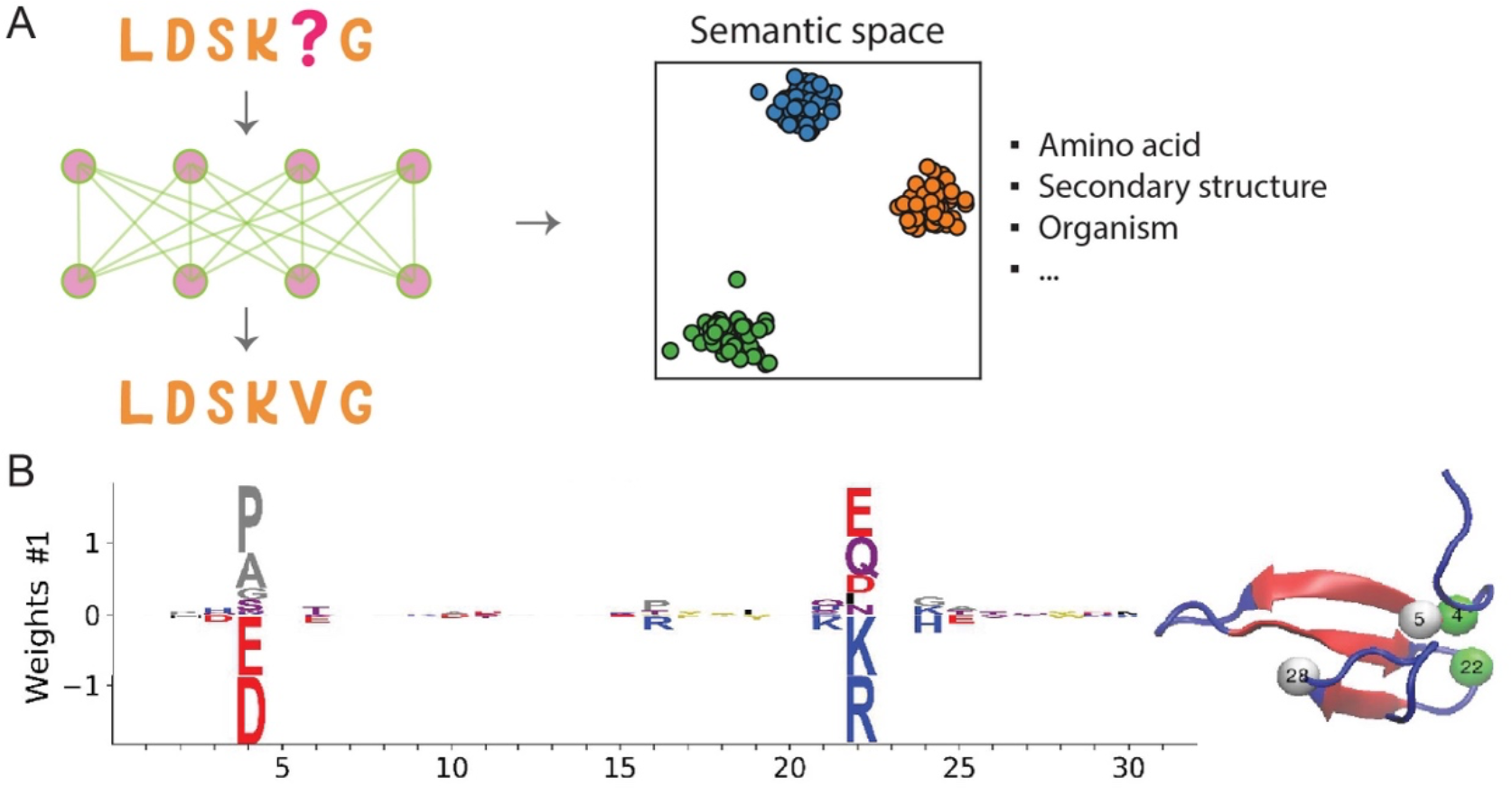
Deciphering protein sequence semantics with generative models. (A) Language models embody the semantics encoded in sequences, capturing various elements including amino acid types, secondary structure, and information at the organism level, to name a few. (B) The weight logo associated with a hidden unit of the RBM model imparts biologically meaningful information. For example, the interactions of charged residues between residues 4 and 22 are underscored. This figure, adapted from Ref (72), is distributed under the terms of the Creative Commons Attribution License.

Hie et al. also utilized an alternative language framework to examine the semantics and fitness of viral proteins (73). The language model’s sequence embedding captures the semantics, where sequences in different semantic landscape areas may suggest antigenic differences. Furthermore, sequences with high probability in the language model indicate increased fitness. These two factors collectively influence the likelihood of a new viral protein mutation evading detection by the immune system.

Although language models have shown great success in learning the semantics of natural language, their main focus has been on text-based inputs. However, protein sequences involve more than just text-based information, and also encompass evolutionary, biological and physical priors that traditional language models maybe unable to capture well (18, 74, 75). As a result, it is necessary to develop new language models that can incorporate these priors and go beyond superficial semantic visualization in order to better understand protein sequences.

Besides language models, using a restricted Boltzmann machine, a generative model akin to the MaxEnt model, Tubiana et al. managed to extract meaningful biological features from the learned representation (72). Certain hidden units were ascribed biological relevance, such as a hidden unit trained with the Kunitz Domain MSA that represented the interaction between charged residues at two separate sites (**Fig 6B**). While it is unclear if each hidden unit can be assigned a definite biological meaning, this analysis offers valuable insights. This technique is somewhat related to predicting protein contacts using evolutionary information and protein sector analysis (41, 63).

### Applications in Enzyme Engineering

Generative models have been employed to design functional enzymes, as summarized in **Table 2**. For example, Russ et al. utilized DCA to design chorismate mutase homologs, resulting in 481 functional sequences from 1618 tested, an impressive success rate of ∼30% (76). Biochemical assays characterized five designed sequences, demonstrating activity similar to their natural counterparts. Repecka et al., using GANs, investigated malate dehydrogenase and found activity in 13 of 55 tested sequences (77). Employing VAEs, Hawkins-Hooker et al. examined bacterial luciferase luxA, with nine out of 11 tested sequences exhibiting activity (78). Giessel, also utilizing VAEs, explored ornithine transcarbamylase (79). Protein language models, in addition to models trained on MSA, have also been used to create lysozyme variants (80).

**Table 2.** Application of generative AI in enzyme engineering.

| Enzyme | Generative Model | Mutant Type | Designs Functional? | Improved Activity vs. Wild-type? |
| --- | --- | --- | --- | --- |
| Chorismate Mutase | MaxEnt (76) | Multiple | ✓ |  |
| Malate Dehydrogenase | GAN (77) | Multiple | ✓ |  |
| Luciferase LuxA | VAE (78) | Multiple | ✓ |  |
| Lysozyme | Language (80) | Multiple | ✓ |  |
| Ornithine Transcarbamylase | VAE (79) | Multiple | ✓ | ✓ (2.3 fold) |
| Luciferase RLuc | MaxEnt (64) | Single | ✓ | ✓ (2.0 fold) |

In most of these enzyme engineering endeavors, the tested sequences do not surpass wild-type or natural enzymes in performance, and even when improvements occur, numerous mutations has to be introduced. While such diverse variants with natural-like functions highlight the generative potential of machine learning models, their practical applications face limitations due to factors such as increased cost, complexity, potential ethical concerns, and poor solubility in some cases (81). Hence, there is a need to develop innovative methods to identify beneficial variants, a task that falls within the realm of rational enzyme engineering.

Rational enzyme engineering demands a deeper understanding of enzyme action. As discussed in a previous section, the evolutionary information in various enzyme components is linked to distinct physicochemical properties, specifically, active sites and enzyme scaffolds are primarily related to the evolutionary pressure of enzyme activity and stability, respectively (60). Recently, we applied this concept to RLuc luciferase (64). The evolution-catalysis relationship, as demonstrated, still holds true for RLuc when evaluated using current biochemical data. We then follow the relationship to design mutants with potentially enhanced properties. Four out of eight single mutants surrounding the substrate exhibited improved activity compared to the wild-type, with minimal changes in stability. In contrast, three out of six expressed single mutants in the enzyme scaffold displayed increased stability but reduced activity. The overall experimental results confirms the evolution-catalysis relationship. Intriguingly, the maximum fold enhancement in activity achieved through a single mutation in RLuc is 2.0, which is on par with the best generative design resulting from multiple mutations (on average eight mutations) in ornithine transcarbamylase, with a fold increase of 2.3 (79). The results highlights the potential of using generative modeling for enzyme sequences in rational enzyme engineering.

### Summary and Perspective

Generative models, which learn patterns underlying naturally evolved sequences, hold the potential to revolutionize our comprehension of enzymes and ultimately aid in designing efficient biocatalysts. In this review, we touched upon a few topics that focus on generative modeling of enzyme sequences. For a broader interest in machine learning in protein studies, we recommend readers explore other reviews (12–26).

Implementing generative models to investigate enzyme physicochemical properties necessitates a large quantity of both sequencing data and enzyme biochemistry data. Fortunately, metagenomic databases, which house genetic material from microbial communities, are experiencing rapid growth (82). Improvements in sequencing technologies could further boost data collection. However, the slow accumulation of physicochemical data for enzyme variants obstructs the establishment of relationships between sequences and functions. The standardization of enzymatic data reporting is crucial (83), and the application of microfluidics for high-throughput measurement of enzymatic data seems promising (43, 84).

In addition to data, a crucial aspect of machine learning involves selecting a model that aligns with the inductive bias of a specific problem. Many existing models introduced here have been directly borrowed from other disciplines, which may not provide the optimal approach for addressing biological problems. This situation can hinder the ability to learn and extract genuine biological principles. From another perspective, it is worth considering the development of models specifically tailored to the unique requirements of biological research, including, proposing biologically meaningful tokenization, structure-informed sequence embedding, designing good measure for sequence representation quality, etc. (18, 75, 85) Furthermore, interdisciplinary collaboration between experts in enzymology, machine learning, and physics-based models could foster the development of novel models that are better suited for biological problems and can effectively capture the inherent characteristics of biological systems.

While the focal point of our review is the application of generative modeling to enzyme sequences, we acknowledge the transformative potential of applying generative models to protein structures in a bid to enhance enzyme design. Recent breakthroughs in protein structure prediction have presented potent tools for translating protein sequences into structures (86), sparking the development of novel strategies that utilize generative models for de novo protein design. Grounded in the principles of nonequilibrium thermodynamics, generative diffusion models enable protein design under a variety of circumstances, including the scaffold of enzyme active sites (87, 88). However, the endeavor of constructing efficient enzymes de novo remains a challenging task due to many mysteries surrounding enzyme catalysis (89). As a result, physics-based models need to be improved and integrated with generative designs.

Additionally, natural language processing using generative language models, such as ChatGPT (30), can revolutionize our comprehension of enzymes. Specifically, ChatGPT can aid in extracting knowledge from previous enzyme research, expediting the discovery process. GPT-4, for example, envisions a wider range of applications for generative models in enzyme studies than our review outlines (see SI). While some assertions might seem early, the prospects for upcoming advancements are bright.

In summary, the integration of generative models into enzyme studies holds the promise of expanding our knowledge of enzymology and facilitating the development of efficient enzymes for diverse applications, potentially giving rise to a novel domain called “evolutionary catalysis.” To this end, embedding domain expertise into generative models can enhance their capacity to address the specific complexities inherent in enzyme research and engineering.

## Supplementary data

Supplementary data are available.

## Acknowledgments

We thank Yousong Ding and Zhenghan Liao for insightful discussions. One piece of content from ChatGPT was included in the SI.

## Funding

A.W. is supported by the National Institutes of Health R35 GM122472 and the National Science Foundation Grant MCB 1707167. W.J.X. is supported by the startup funding from University of Florida.

## Conflict of interest statement

None declared.

## Notes

### Competing Interest Statement

The authors have declared no competing interest.

