## Supplementary material for "Harnessing Generative AI to Decode Enzyme Catalysis and Evolution for Enhanced Engineering": SI

**ChatGPT's Perspective on Utilizing Generative Models in Enzyme Studies**

*Prompt: How might generative models be applied to the engineering and study of enzymes?*

GPT-4 generates:

Generative models, like those based on the GPT architecture, can be applied to the engineering and study of enzymes in various ways. These models have the potential to accelerate enzyme research, improve our understanding of enzyme function, and facilitate the design of novel enzymes for specific applications. Some applications include:

Protein sequence generation: Generative models can be trained to create new protein sequences based on known sequences in a given enzyme family. These sequences can be used to explore potential enzyme variants with novel functions, increased stability, or improved catalytic activity.

Protein structure prediction: Generative models can be employed to predict the 3D structure of enzymes based on their amino acid sequences. Accurate structure prediction can provide insights into enzyme mechanisms, substrate binding sites, and help guide the design of enzyme inhibitors or activators for therapeutic purposes.

De novo enzyme design: Using generative models, researchers can design entirely new enzymes with custom functions or activities. By incorporating principles of enzyme catalysis and substrate specificity, these models can generate sequences that encode for proteins with desired properties, paving the way for tailored biocatalysts and therapeutics.

Enzyme engineering for improved function: Generative models can be used to suggest mutations or modifications to existing enzymes to enhance their catalytic efficiency, stability, or specificity. This can be particularly useful for industrial applications, where enzymes are often used as biocatalysts in the production of chemicals, pharmaceuticals, and biofuels.

Enzyme-Substrate interaction prediction: Generative models can be employed to predict enzyme-substrate interactions, which can help researchers understand how enzymes recognize and process their substrates. This information can be used to design more efficient enzymes or develop inhibitors that specifically target enzyme-substrate interactions.

Metabolic pathway engineering: By generating enzyme variants with altered activities or specificities, generative models can aid in the design of engineered metabolic pathways for the production of valuable chemicals or degradation of environmental pollutants.

Machine learning-guided directed evolution: Generative models can guide the selection of mutations for directed evolution experiments, which are used to evolve enzymes with improved properties. By intelligently proposing mutations, these models can potentially reduce the number of experimental iterations needed to achieve the desired enzyme characteristics.

Education and training: Generative models can be used to create educational materials, such as tutorials and interactive tools, to help students and researchers understand enzyme structure, function, and engineering principles.

Overall, the application of generative models in enzyme engineering and study holds immense promise for advancing our understanding of enzymatic processes and the development of novel enzymes for various applications.
